# DeepFlu: a deep learning approach for forecasting symptomatic influenza A infection based on pre-exposure gene expression

**DOI:** 10.1101/2020.12.02.407940

**Authors:** Anna Zan, Zhong-Ru Xie, Yi-Chen Hsu, Yu-Hao Chen, Tsung-Hsien Lin, Yong-Shan Chang, Kuan Y. Chang

## Abstract

**Background and Objective:** Not everyone gets sick after an exposure to influenza A viruses (IAV). Although KLRD1 has been identified as a potential biomarker for influenza susceptibility, it remains unclear whether forecasting symptomatic flu infection based on pre-exposure host gene expression might be possible.

**Method:** To examine this hypothesis, we developed DeepFlu using the state-of-the-art deep learning approach on the human gene expression data infected with IAV subtype H1N1 or H3N2 viruses to forecast who would catch the flu prior to an exposure to IAV.

**Results:** The results indicated that such forecast is possible and, in other words, gene expression could reflect the strength of host immunity. In the leave-one-person-out cross-validation, DeepFlu based on deep neural network outperformed the models using convolutional neural network, random forest, or support vector machine, achieving 70.0% accuracy, 0.787 AUROC, and 0.758 AUPR for H1N1 and 73.8% accuracy, 0.847 AUROC, and 0.901 AUPR for H3N2. In the external validation, DeepFlu also reached 71.4% accuracy, 0.700 AUROC, and 0.723 AUPR for H1N1 and 73.5% accuracy, 0.732 AUROC, and 0.749 AUPR for H3N2, surpassing the KLRD1 biomarker. In addition, DeepFlu which was trained only by pre-exposure data worked the best than by other time spans and mixed training data of H1N1 and H3N2 did not necessarily enhance prediction. DeepFlu is available at https://github.com/ntou-compbio/DeepFlu.

**Conclusions:** DeepFlu is a prognostic tool that can moderately recognize individuals susceptible to the flu and may help prevent the spread of IAV.

## 1. Introduction

Influenza is a highly contagious and deadly viral respiratory disease second only to coronavirus disease 2019 (covid-19). There are about 3 to 5 million severe cases of influenza worldwide every year, resulting in approximately 291,000 to 646,000 deaths annually [1, 2]. Influenza A caused by influenza A virus (IAV) accounts for about 75% of all influenza infections [3]. IAV is also responsible for 1918, 1957, 1968, and 2009 influenza pandemics [4]. About 2.3% of the adults have contracted the IAV [5, 6], but not everyone gets sick after an exposure to IAV [7]. Slightly more than half of young adults would get infected and develop symptoms of acute upper respiratory tract infection after a controlled IAV exposure; the rest remain either asymptomatic or uninfected [8, 9].

Due to the huge socioeconomic and health burden of influenza [10, 11, 12], it is important to identify and anticipate who will contract influenza and become ill. Delaying the recognition of infected persons with IAV will increase the viral transmission [13] and sometimes worsen the infection, thereby increasing the death toll. Although vaccination is recommended to prevent the influenza and improve public health in schools, hospitals, communities, and societies [14], flu vaccines are not necessarily working for everyone [15]. Instead, foreseeing who will succumb to IAV infection prior to viral exposure could be an alternative to prevent the spread of IAV and reduce the death toll from the IAV and its related complications.

It remains unclear whether host susceptibility to IAV is predictable based on pre-viral-exposure host gene expression. KLRD1 expression has been recently identified as the inverse biomarker for predicting influeuza susceptibility based on a meta analysis [16]. Besides, two prediction models have shown promising results for forecasting host susceptibility to respiratory syncytial virus (RSV) [9]. One is an epsilon support vector regression model [17] with a radial basis function kernel [9] learning directly from host gene expression. The other is a LASSO regularized regression model [9] based on modulation of biological pathways, deriving from the Hallmark gene sets [18]of the Molecular Signature Database (MSigDB). However, no one has yet demonstrated whether it is feasible using any advanced machine learning approach to forecast the onset of IAV infection prior to viral exposure, and if so, whether this forecast can outperform the KLRD1 biomarker.

Forecasting host susceptibility to IAV differs from detecting IAV infection during the onset of influenza symptoms based on host gene expression. Although both host gene expression responses are influenced by the same factors such as host genetics [19], immunological memory [20], seasonal and circadian changes [21], age [22], and sex [23, 24] they attend to different times. Forecasting IAV susceptibility centers on the time prior to an IAV exposure, but IAV-infected hosts can be detectable only after exposure to the IAV [8, 23, 25].

It has been a success to detect IAV infection based on host gene expression. The peripheral blood gene expression signatures of post IAV infection different from those of bacterial and other respiratory viral infections can be used to detect symptomatic IAV infection [8, 23], which have been applied to the real-world cases of 2009 H1N1 pandemic with over 90% accuracy. In fact, comparing top 50 discriminative genes [25], the infection signatures across different IAV substrains are so similar [25] that some have exhibited cross-strain detectability [26]. Remarkably, a multi-cohort analysis further demonstrates that detecting symptomatic subjects infected with IAVs can count on only 11 genes derived from an influenza meta-signature [23]. Whether such success can occur in forecasting host susceptibility to IAV needs to be unraveled.

To our knowledge, this is the first study to attempt the-state-of-the-arts deep learning to forecast host susceptibility to IAV prior to a viral exposure. In this study, we report the development of a deep learning model called DeepFlu.

## 2. Methods

### 2.1. Datasets Description

The human peripheral blood gene expression datasets from the NCBI database Gene Expression Omnibus (GEO) were used. Each dataset was standardized by RMA normalization [27]. GEO GSE52428 [25] which was done by the Affymetrix Human Genome U133A 2.0 array with 22,277 probes was the primary dataset for training and testing. 24 and 17 human subjects who aged 19 to 41 were inoculated intranasally with influenza A H1N1 (A/Brisbane/59/2007) or H3N2 (A/Wisconsin/67/2005), respectively. All subjects were healthy prior to the inoculation and monitored about every 8 hours over the course of 7 days. Consequently, 9 of the 24 H1N1 subjects and 8 of the 17 H3N2 subjects became symptomatic IAV-infected. 3 H1N1 subjects who developed symptoms but not infected by the H1N1 viruses were not classified as symptomatic IAV-infected. The rest were either asymptomatic or uninfected. Consequently, a total of 382 H1N1 and 267 H3N2 expression records were collected (Fig 1).

**Figure 1:**
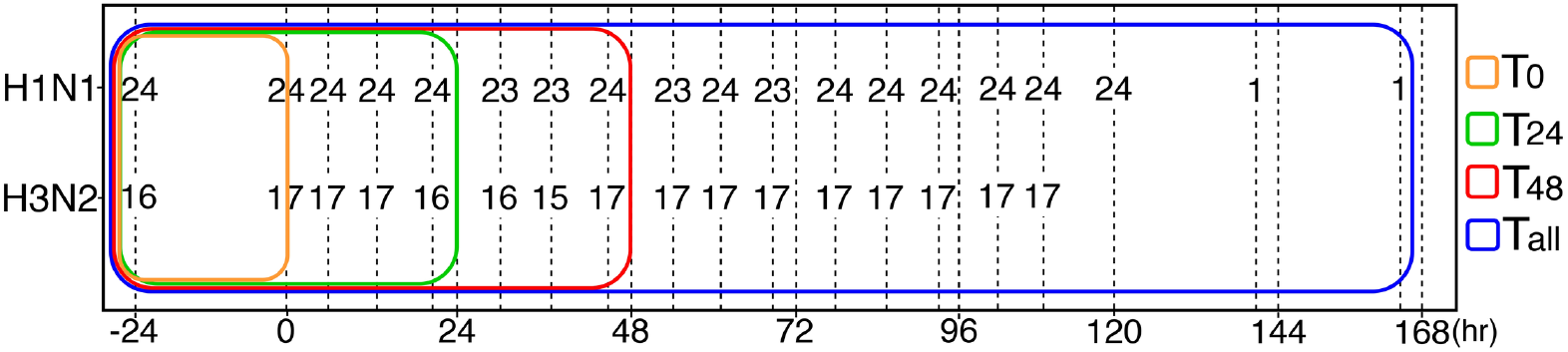
Datasets in four different time spans. T_0_, T_24_, T_48_, and T_all_ represent the data taking from 1 day before influenza exposure up to immediately before exposure, 24h, 48h, and 168h after exposure, respectively. Each number in the dash line is the number of subjects at that time point.

The external validation set came from GEO GSE73072 [28]. The experiments of DEE2 and DEE3 in this dataset were the same as those of GSE52428, but done by the Affymetrix Human Genome U133A 2.0 array with 12,023 genes instead. Thus, the external validation on additional subjects were DEE4 and DEE5, where 6 of the 19 H1N1 subjects and 13 of the 21 H3N2 subjects were symptomatic infected.

Being symptomatic IAV-infected is defined as the presence of both clinical flu symptoms and microbiological infection. Technically speaking, a symptomatic infected subject should hold the modified Jackson score ≥ 6 for at least 5 consecutive days and a positive viral culture or RT-PCR viral test [29] for at least 2 consecutive days. The modified Jackson score [30] assesses each of eight symptoms on a severity scale of 0 to 3, including sneeze, cough, malaise, runny nose, stuffy nose, sore throat, myalgia, and headache, to determine the presence or absence of flu symptoms. The viral culture or RT-PCR test is used to confirm the existence of IAV infection. Once the subject was identified as symptomatic IAV-infected, all of the subject’s records were marked accordingly.

### 2.2. Leave-one-person-out cross-validation

Due to the limited gene expression data, a leave-one-person-out (L1PO) cross-validation strategy was applied to the primary dataset for model evaluation. The L1PO crossvalidation which evaluates how well a model predicts the outcome of an unknown subject upon to viral exposure–who would become symptomatic infected or not–involves a rotation of all individuals partitioning into one-person validation set and the remaining set. Models which were trained using the observations of the remaining set were tested on the validation set. Because of stochastic nature of the models, we repeated the procedure 100 times to estimate the overall model performance.

### 2.3. DeepFlu Layout

Fig 2 illustrates the architecture of DeepFlu designed for predicting symptomatic IAV infection. DeepFlu is a Deep Neural Network (DNN), also known as multilayer perceptron, which is a feedforward neural network that can distinguish linear non-separable patterns [31]. A series of model assessment was taken (See the supplementary material). DeepFlu is a 6-layer DNN, consisting four fully connected hidden layers each with 100 nodes, one input layer to extract the gene expression information, and one output layer to predict the presence of symptomatic flu infection.

**Figure 2:**
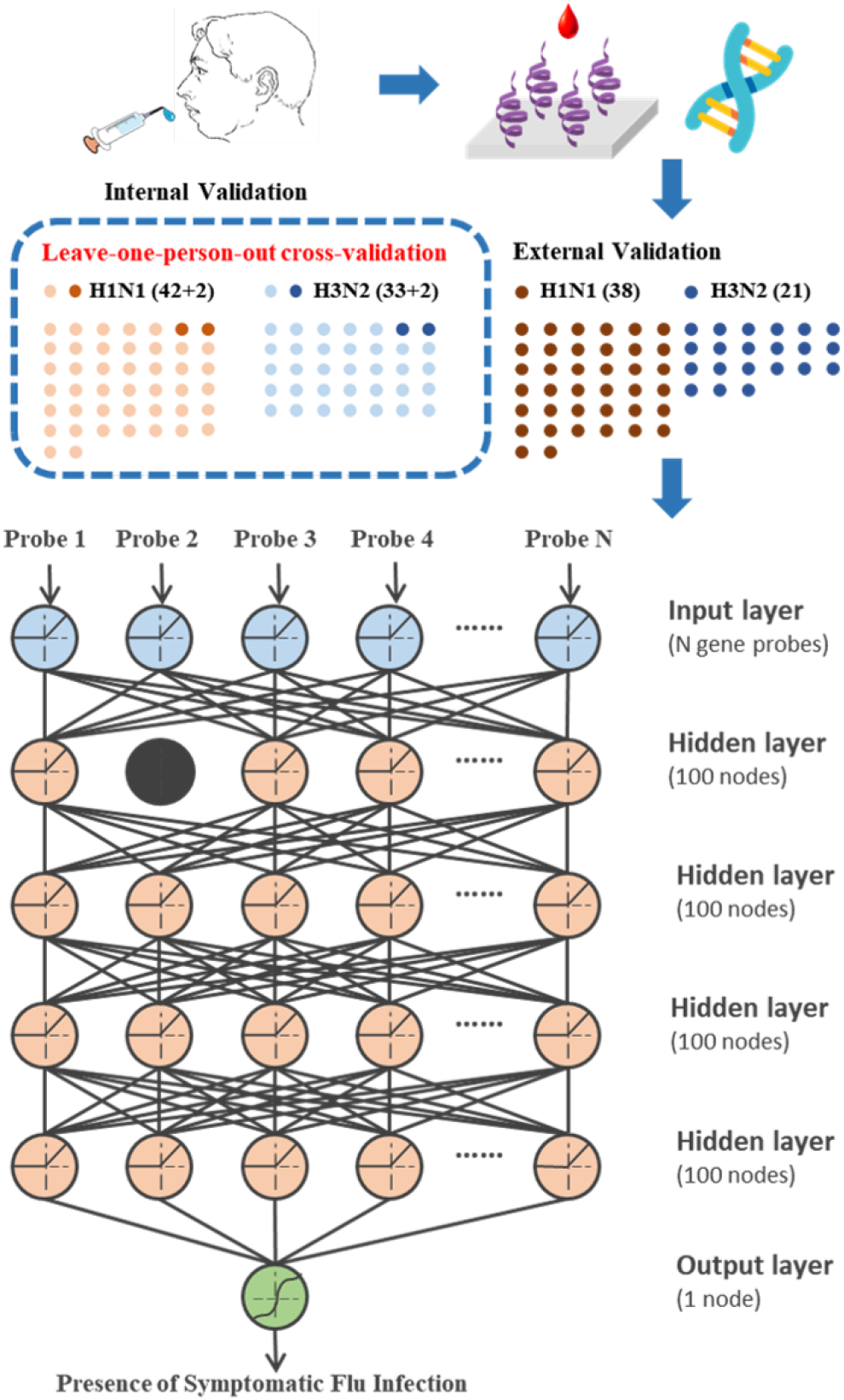
DeepFlu Architecture. DeepFlu is a 6-layer DNN. Its input layer consists of *N* nodes corresponding to gene expression probes. Each of the four fully connected hidden layers contains 100 nodes, where the black node represents the dropout. The output layer utilizes single node to generate a binary outcome.

The following parameters and methods were used in DeepFlu. Both the input and hidden layers used a rectified linear unit (ReLU) function [32] as the activation function, for it could overcome the vanishing gradient for nonnegative inputs. The output layer transformed a logistic regression with a sigmoid function, whose outcome was strictly increasing and [0, 1], into a binary decision. To minimize the loss measured by binary cross-entropy, adaptive moment estimation (ADAM) optimization function [33], which used the moving averages of the past gradient and squared gradient to update model weights, was applied during backpropagation, for ADAM could work effectively for high-dimensional parameter spaces. In addition, 150 training epochs were run with a learning rate of 0.001 without weight decay and 10% dropout as regularization was applied to the first hidden layer to avoid overfitting. The performance of DeepFlu was evaluated by both L1PO cross-validation on the primary dataset and the external validation.

### 2.4. Convolutional Neural Network

Convolutional neural networks (CNNs) [34] are one of the most popular deep learning models, especially for image recognition [35, 36]. Many well-known image recognition models such as AlexNet [37] and ResNet [38] are CNNs. However, it remains unclear how CNNs perform in gene expression data.

We built a 10-layer CNN model, extending from the 6-layer DeepFlu by adding two convolutional layers and two pooling layers. CNNs, which can acquire spatial relations of data, can be divided into two sections. The first section is to extract features through alternate convolutional filters and pooling layers, and the second section is to learn through fully connected layers. Moreover, the activation function of the convolutional layers was a ReLU function [32] and the filter of the pooling layers was a max pooling function. The rest of parameters and methods used in the CNN model were adopted from those of DeepFlu. Because the CNN model had more layers and parameters, the CNN model would have more epochs than DeepFlu. The CNN models can be described simply as follows: the input is a tensor *X* ∈ **R**^*w×h×d*^, where *w, h*, and *d* are the image width, height, and depth, respectively, and the depth refers to the number of channels. The convolution transforms the input *X* using *m* filter tensors *K* ∈ **R**^*n×n×d*^ with *q* zero-paddings and *s* strides into *m* feature maps 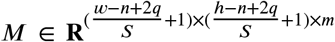, where *n* is the order of filter kernels, *d* keeps the same number of channels as the input, and *m* which determines the number of the output channels is the number of filter kernels. To effectively reduce the dimensions of feature maps, the pooling layer adopts a filter kernel. The pooling layer transforms the feature map using a pooling filter tensor *P* ∈ **R**^*q×q×m*^ into a pooled feature map, where *q* is the order of pooling kernels and *m* is the channel number of feature maps. Excluding the convolutional and pooling layers, the makeup of the CNN models used in this study is the same as that of DeepFlu.

### 2.5. Random Forest

Random Forest (RF) is a classification method using an ensemble of bagged decision trees with randomly selected features [39]. The RF algorithm embeds the concept of random subset feature selection into bagged decision trees which are trained on random samples with replacement to avoid overfitting. To determine the class of the predicted case, each decision tree which is best split by a small fraction of randomly selected features casts a vote independently under majority rule. In this work, the parameters and methods used for the RF models were adopted from the Chang study [40]. Each RF classifier had 1,000 decision trees. Gini impurity [41] was used to measure the quality of a split by a selected feature and when looking for a best split, the number of tried features was set to be square root of the total features [42].

### 2.6. Support Vector Machine

SVM is also a classification method that searches an optimal hyperplane to separate two classes with maximum margin [43]. In this work, a radial basis function kernel [44] was used for our SVM models, whose cost and gamma parameters [17] for the kernel were set as 1 and one over the number of features.

### 2.7. Performance Comparison

The performances of the predictive models were measured by sensitivity, specificity, precision, accuracy, area under receiver operating characteristic curve (ROC), and area under precision recall (PR) curve. These evaluation measurements were calculated as follows:

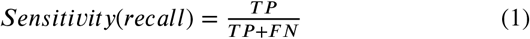

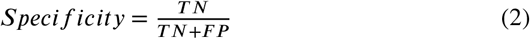

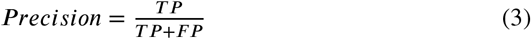

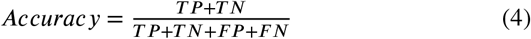

where TP, TN, FP, and FN represent true positives, true negatives, false positives, and false negative, respectively. Sensitivity, specificity, precision, and accuracy are the percentage of the correct predictions on positive data, negative data, predicted positive data, and all the data, respectively.

Like accuracy, ROC and PRC are used to evaluate the overall performance of a binary classifier [45, 46]. ROC and PRC are based on dynamic positive-negative prediction thresholds instead of a static threshold like accuracy. ROC is the area under the curve which plots true positive rates (sensitivity) against false positive rates using different prediction thresholds; similar to ROC, PRC is the area under the curve by plotting precision against recall rates (sensitivity). ROC and PRC are in the range [0,1]. The higher ROC or PRC, the better the classifier. A perfect classifier has ROC or PRC equal to 1. However, unlike ROC, PRC only focuses on positive cases. The ROC value tends to be overly optimistic for an imbalance dataset, but PRC does not [47].

## 3. Results

### 3.1. Best Learning Method for the Gene Expression Data

By randomly dividing the influenza gene expression data into 80% training and 20% testing data, Table 1 shows the performances of four different predictive modeling approaches using the full 22,277 array probes. The setting of the training and testing data used here, which were not exclusive to those prior to exposure, differed from most of the experiments throughout this study. All the four models, CNN, DNN, RF, and SVM, demonstrate that the special gene expression signals of the flu infected subjects could be learned. Among the four models, DNN generally performed the best, reaching averagely 96.1% accuracy, 0.957 AUROC, and 0.963 AUPR for H1N1; 95.4% accuracy, 0.954 AUROC, and 0.965 AUPR for H3N2. Thus, DNN was chosen as the backbone of DeepFlu for the following development.

**Table 1:** Model comparison for distinguishing symptomatic flu-infected subjects

| Method | Accuracy | Sensitivity | Specificity | Precision | AUROC | AUPR |
| --- | --- | --- | --- | --- | --- | --- |
| **H1N1** |  |  |  |  |  |  |
| CNN | 0.955 | **0.951** | 0.960 | 0.929 | 0.955 | 0.949 |
| DNN | **0.961** | 0.936 | **0.978** | **0.964** | **0.957** | 0.963 |
| RF | 0.911 | 0.835 | 0.960 | 0.927 | 0.898 | **0.969** |
| SVM | 0.841 | 0.639 | 0.891 | 0.881 | 0.789 | 0.859 |
| **H3N2** |  |  |  |  |  |  |
| CNN | 0.937 | 0.962 | 0.914 | 0.940 | 0.946 | 0.951 |
| DNN | **0.954** | **0.975** | **0.932** | **0.942** | **0.954** | 0.965 |
| RF | 0.924 | 0.930 | 0.921 | 0.927 | 0.925 | **0.984** |
| SVM | 0.867 | 0.851 | 0.888 | 0.898 | 0.866 | 0.913 |

AUROC: area under receiver operating characteristic curve; AUPR: area under precision-recall curve; CNN: convolutional neural network; DNN: deep neural network; RF: random forest; SVM: support vector machine

### 3.2. Time Spans on Prediction Performance

Based on the L1PO cross-validation, Table 2 compares the prediction performances of DeepFlu trained under different time spans. T_0_, T_24_, T_48_ and T_all_ represent the expression data taking from the beginning (one day before exposure) to the moment of exposure, one day after exposure, two days after exposure, and one week after exposure, respectively (Fig. 1). In other words, T_0_ contains only the expression data without viral exposure. The results show that training by the data prior to viral exposure (T_0_) was the best for DeepFlu to predict whether one would catch the flu and develop symptoms after an exposure, achieving 70.0% accuracy, 0.787 AUROC, and 0.758 AUPR for H1N1 and 73.8% accuracy, 0.847 AUROC, and 0.901 AUPR for H3N2 (Fig. 3A-B).

**Table 2:** Performance comparison of DeepFlu trained under different time spans by L1PO cross-validation

| Span | Accuracy | Sensitivity | Specificity | Precision | AUROC | AUPR |
| --- | --- | --- | --- | --- | --- | --- |
| **H1N1** |  |  |  |  |  |  |
| $T_{All}$ | 0.637 | 0.556 | 0.668 | 0.618 | 0.661 | 0.638 |
| $T_{48}$ | 0.658 | 0.602 | 0.689 | 0.662 | 0.713 | 0.701 |
| $T_{24}$ | 0.669 | 0.613 | 0.715 | 0.679 | 0.725 | 0.712 |
| $T_0$ | **0.700** | **0.616** | **0.822** | **0.718** | **0.787** | **0.758** |
| **H3N2** |  |  |  |  |  |  |
| $T_{All}$ | 0.638 | 0.707 | 0.548 | 0.649 | 0.668 | 0.688 |
| $T_{48}$ | 0.661 | 0.683 | 0.635 | 0.688 | 0.679 | 0.744 |
| $T_{24}$ | 0.689 | 0.689 | 0.723 | 0.759 | 0.759 | 0.806 |
| $T_0$ | **0.738** | **0.722** | **0.756** | **0.770** | **0.847** | **0.901** |

AUROC: area under receiver operating characteristic curve; AUPR: area under precision-recall curve; L1PO: leave one person out

**Figure 3:**
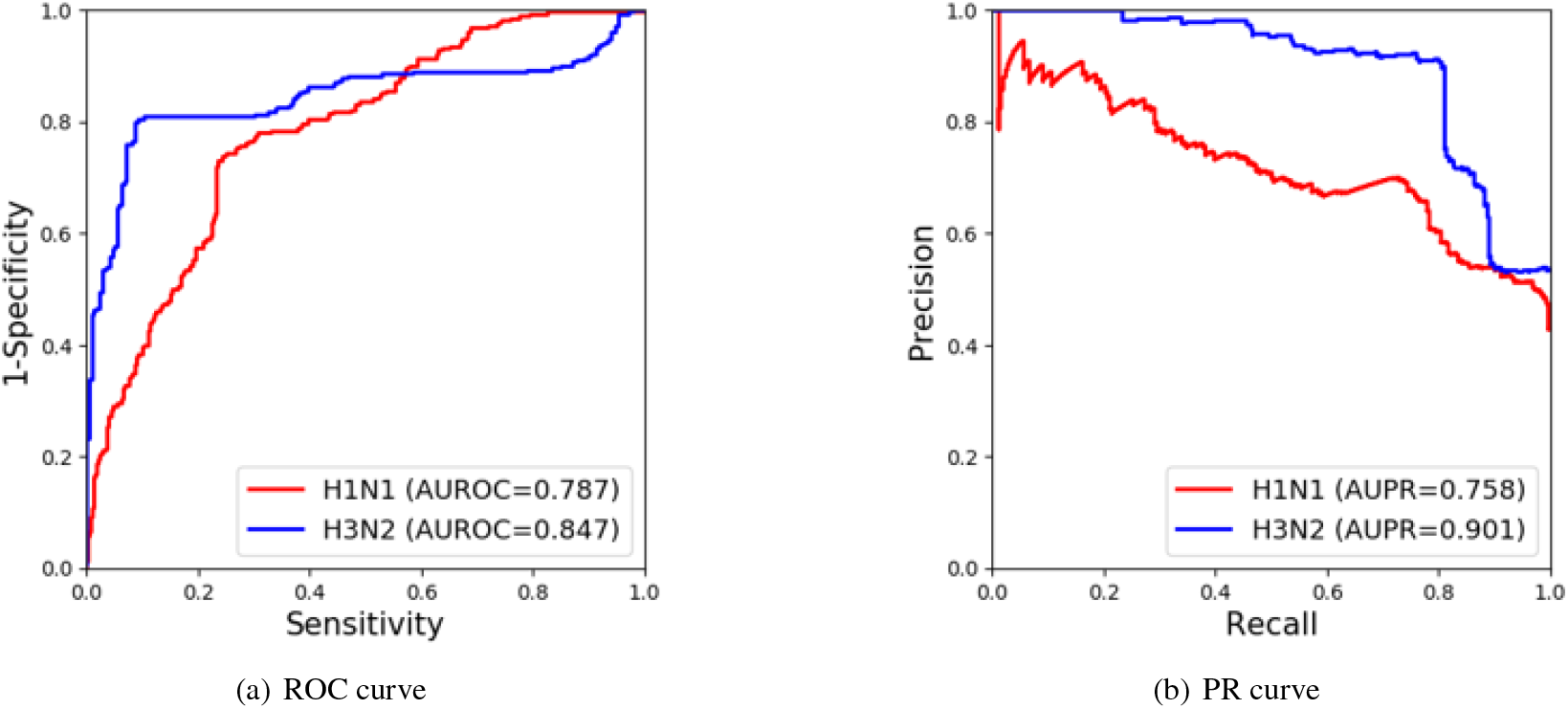
Internal Validation for DeepFlu.

### 3.3. Feature Sets on Prediction Performance

Based on the L1PO cross-validation test focusing on host gene expression prior to exposure (T_0_), Table 3 compares the performances of the four predictive models using three different feature sets for forecasting symptomatic flu-infected subjects. Besides DeepFlu, the three other predictive models were CNN, RF, and SVM; the three feature sets were the complete probe set of the microarray (22,277 features), the Hallmark genes [18] of MSigDB (4,164 corresponding features), and the top discriminative genes for distinguishing IAV post infection (64 and 61 corresponding features for H1N1 and H3N2 respectively) [17]. The results indicate that the prediction models using more features generally performed better and DeepFlu achieved the overall best performance using the full 22,277 features for both H1N1 and H3N2. For H1N1, DeepFlu had 70.0% accuracy, 0.787 AUROC, and 0.758 AUPR; for H3N2, it reached 73.8% accuracy, 0.849 AUROC, and 0.901 AUPR.

**Table 3:**
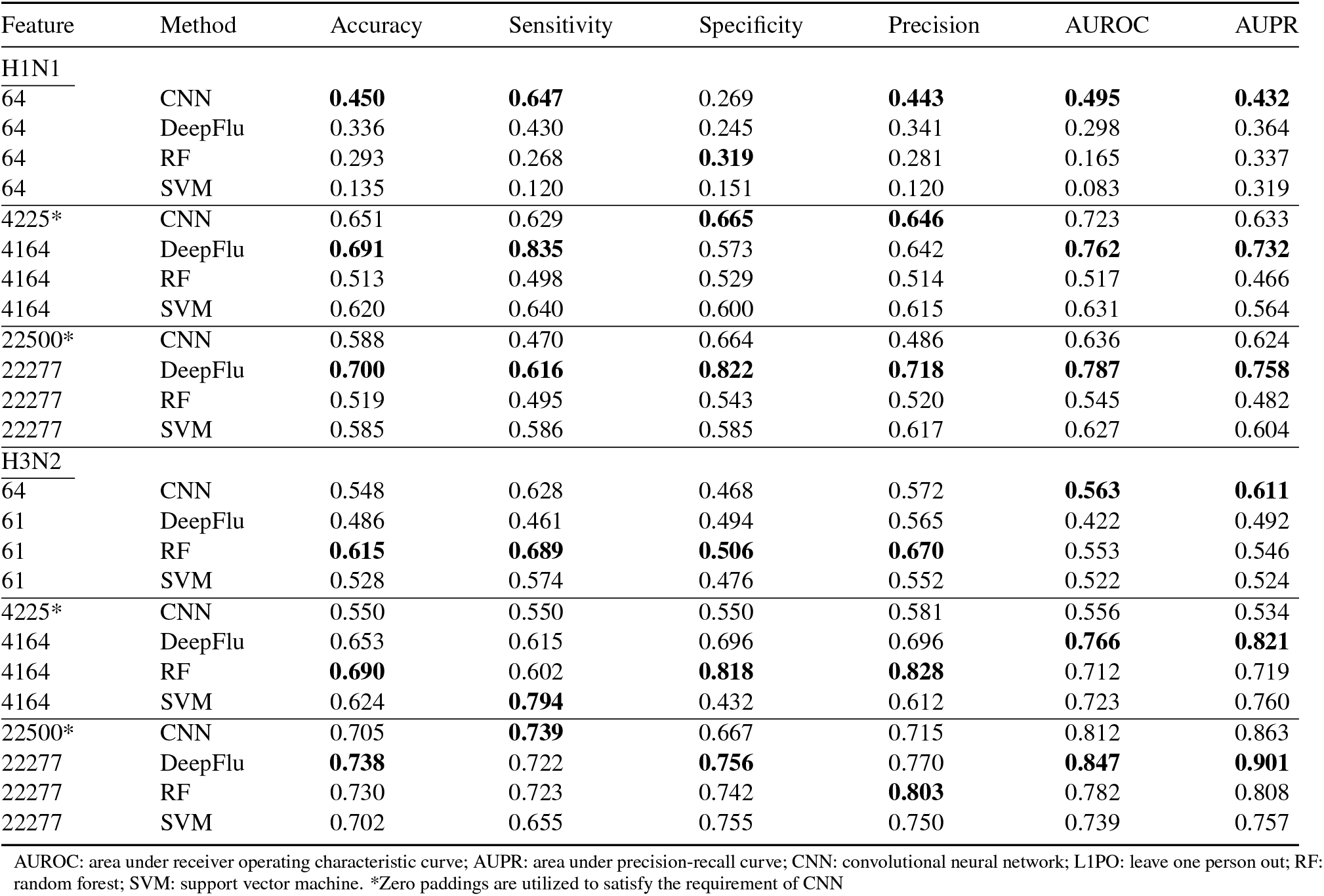
Performance comparison using different feature sets trained under T_0_ by L1PO cross-validation

Compared to using the full 22,277 features, DeepFlu gave a comparable but weaker performance by using 4,164 features. Still it outperformed the other three models.

For using the top discriminative genes of IAV post infection, all the four models performed poorly. All the four models using the limited gene set gave the worst AUROC and AUPR values for H1N1. For H3N2, the models performed slightly better than a random guess.

### 3.4. Mixed Training Data on Prediction Performance

Because the post infection signatures of H1N1 and H3N2 were similar, sharing up to 90% of top 50 discriminative genes [25], and that of H3N2 could be used to detect H1N1 infection [26], it was speculated that extending the training set by mixed data of H1N1 and H3N2 might enhance prediction performance. Instead of training by the expression data for the same IAV strain as seen in Table 3, Table 4 compares the performance of the L1PO cross validation of the predictive models trained by the mixed T0 data of H1N1 and H3N2. Table 4 shows that despite a drop in performance, DeepFlu was still the best for forecasting symptomatic influenza A-infected subjects prior to viral exposure. For H1N1, DeepFlu using the full 22,277 features of the mixed training data achieved 72.1% accuracy, 0.752 AUROC, and 0.676 AUPR, not better than that trained only by the H1N1 data with 70.0% accuracy, 0.787 AUROC, and 0.758 AUPR. Similarly, as for H3N2, DeepFlu using the mixed data dropped performance than using the H3N2 data alone, but retained a high performance, keeping more than 0.790 AUROC and AUPR. The results indicate that the mixed H1N1 and H3N2 training data neither improved identifying who were susceptible to H1N1 nor those to H3N2 infection.

**Table 4:**
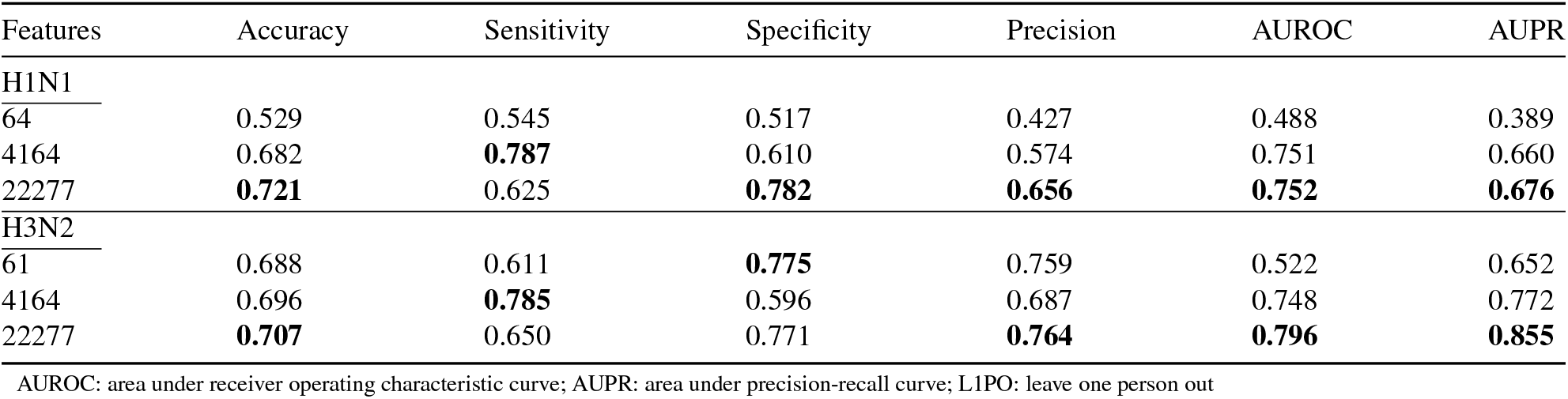
Performance comparison of DeepFlu trained using different feature sets by mixed H1N1 and H3N2 T_0_ data by L1PO cross-validation

### 3.5. Overfitting Diagnosis of Model Parameter

An overfitting diagnosis on DeepFlu was performed using the L1PO cross validation in terms of the number of nodes per hidden layer. The steps for building each DNN model were the same as aforementioned except the number of nodes per hidden layer. To detect overfitting or too many parameters, we looked for when the model may fit the training set very well, but begin to fail to predict for new cases. Figure 4 illustrates the overfitting diagnosis. When the number of nodes per hidden layer was more than 100, the model became overfitting. DeepFlu with 100 nodes per hidden layer demonstrates an optimal performance without overfitting (Fig.4).

**Figure 4:**
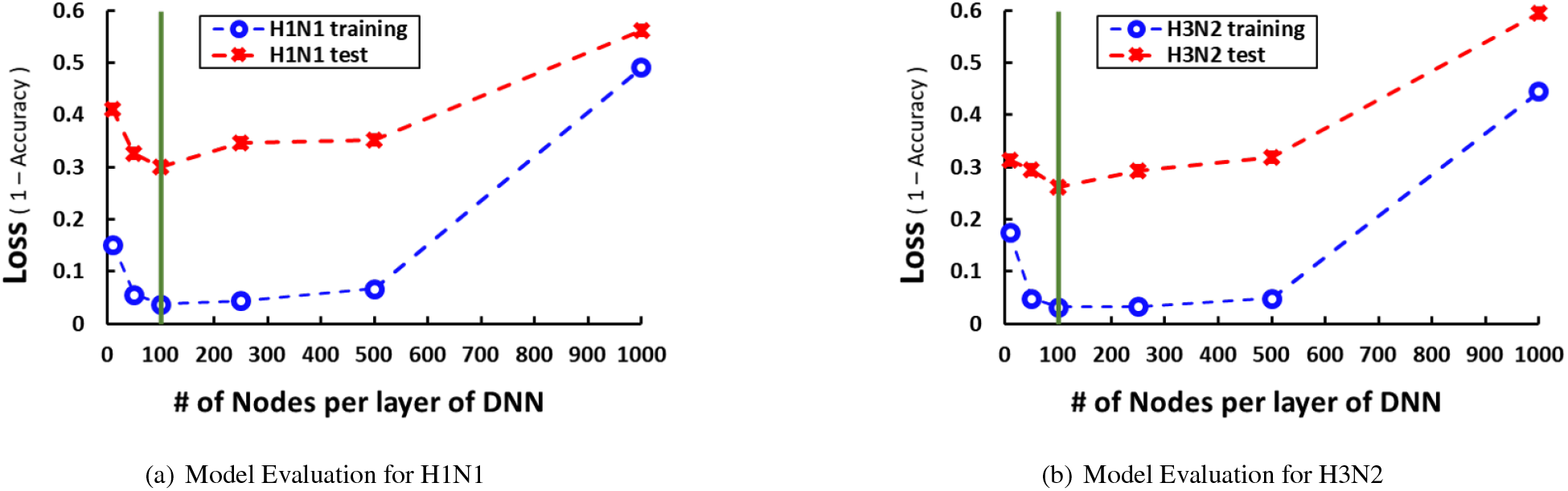
Overfitting Diagnosis.

### 3.6. External Validation on DeepFlu

To examine the robustness of DeepFlu, an external validation of 19 H1N1 and 21 H3N2 subjects were applied. KLRD1, the recently found biomarker of influenza susceptibility, was used for comparison. Table 5 shows the performance of DeepFlu on the external validation. The H3N2 data here came from the same discovery cohort of KLRD1 while none of the validation data were seen by DeepFlu. Regardless, DeepFlu achieved 71.4% accuracy, 0.700 AUROC, and 0.723 AUPR for H1N1; 73.5% accuracy, 0.732 AUROC, and 0.749 AUPR for H3N2, surpassing the KLRD1 biomarker (Fig. 5A-B).

**Table 5:** External validation of DeepFlu

|  | Accuracy | Sensitivity | Specificity | Precision | AUROC | AUPR |
| --- | --- | --- | --- | --- | --- | --- |
| H1N1 | 0.714 | 0.733 | 0.767 | 0.700 | 0.700 | 0.723 |
| H3N2 | 0.735 | 0.714 | 0.750 | 0.667 | 0.732 | 0.749 |

AUROC: area under receiver operating characteristic curve; AUPR: area under precision-recall curve

**Figure 5:**
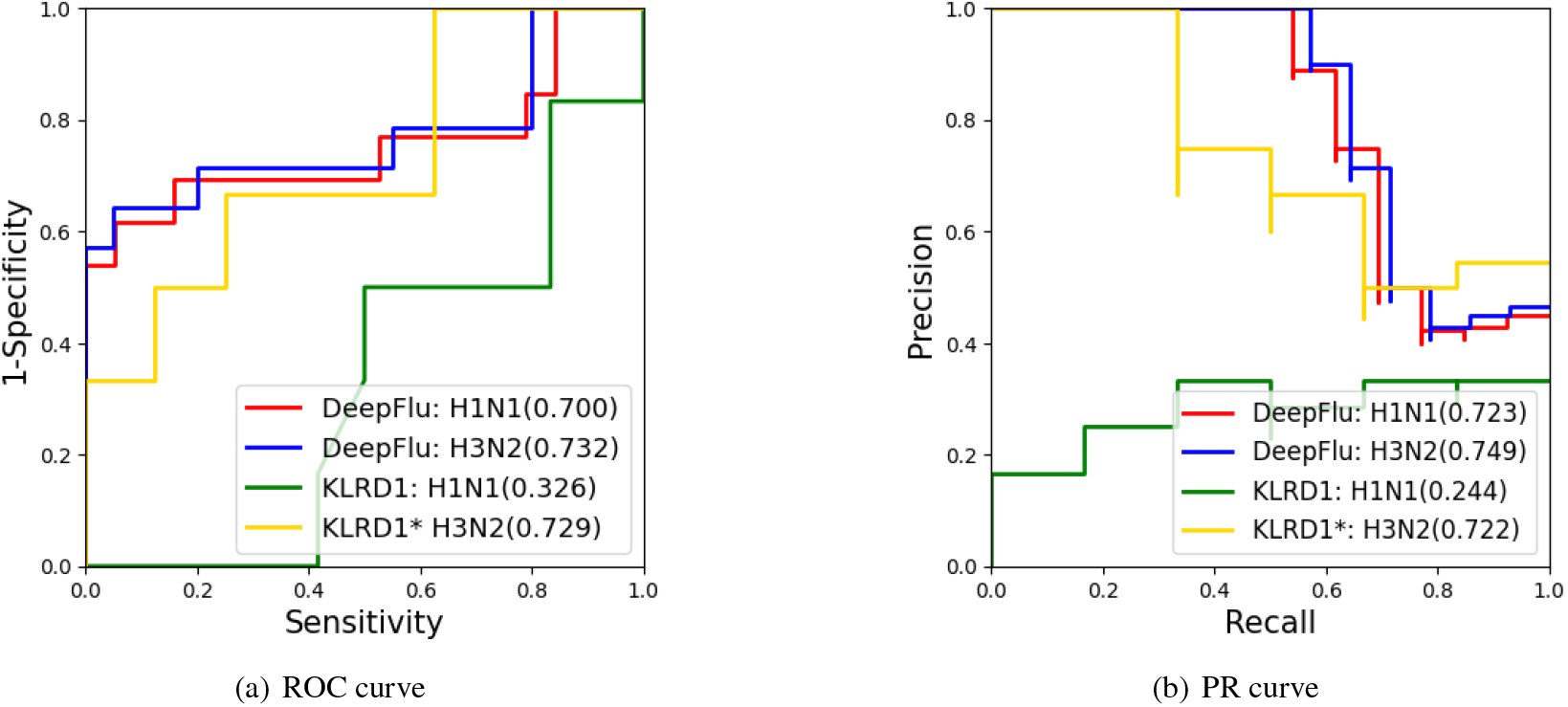
External Validation for DeepFlu. *The H3N2 data here come from the same cohort for discovering the KLRD1 biomarker.

## 4. Discussion

According to the L1PO cross-validation and external validation results, we have demonstrated that it is possible based on pre-IAV-exposure host gene expression to forecast whether an individual would exhibit flu onset immediately after an exposure to IAV by deep learning. The results are consistent with the previous findings [9, 16]. All the studies demonstrate the possibility of forecasting susceptibility of respiratory viral infections.

Deep learning has been demonstrated to work on various data. Although this gene expression data is not like the standard large data, we demonstrate that deep learning can still work. DeepFlu with 100 nodes per hidden layer was not overfitting, but achieved the optimal performance for predicting influenza susceptibility (Fig. 4). Two major reasons why DeepFlu works well on the holdout samples are using dropout, a regularization step, to avoid overfitting and using L1PO cross-validation to select a robust model.

Feature selection can be important for predicting IAV susceptibility. Not only does DeepFlu outperform the KLRD1 biomarker alone, but also our results generally show that the larger the feature size, the better the prediction performance. In fact, DeepFlu using the complete 22,274 features is the optimal model. The DNN model which used the Hallmark genes of MSigDB, about only one-fifth of the complete features, could still reach a comparable performance, indicating that there might exist limited candidate genes closely relating to one’s susceptibility to IAV. However, the underlying biological mechanism might be too complex to learn well with the selected features. Instead entire features became a better alternative.

The top 50 discriminative post-IAV-infection genes was inadequate to be used as features for this problem. The models using the limited IAV infection genes all consistently performed poorly compared with the others. This poor result suggests that the IAV infection genes, which are stimulated after viral exposure, may significantly differ from the genes relating to one’s susceptibility to IAV, which may regulate and control immune responses. In other words, the biological mechanisms behind IAV post infection, such as inflammatory responses [7], might differ from those behind IAV immunity.

The results also indicate that DeepFlu performs better for recognizing gene expression patterns of IAV immunity than the other three models, including the CNN models. DeepFlu outperformed the CNN model in both random setting (80% training and 20% test data) and L1PO setting, perhaps because gene expression profiles did not conform to the CNN properties–neither subsampling gene chips would hold the original meaning nor the same expression patterns could happen in different regions of gene chips.

Why forecasting H3N2 susceptibility could work better than forecasting H1N1 susceptibility is unsettled. We suspect the length of latency period may be a factor, for the maximal symptoms of H3N2 and H1N1 patients peak at day two and day four, respectively [9]. The H3N2 models foresee clearer, perhaps because T_0_ is closer to the symptom onset of H3N2 than that of H1N1.

Moreover, using which time span to train is also important. The results support that the best outcome happened when both the training and test data came from the same time span—prior to exposure (T_0_). Having other time spans might bring in other signals or noises, thus lowering the performance.

The performance of forecasting may still have room to improve. However, by mixing H1N1 and H3N2 records to expend the training data may not be the way to enhance prediction performance. Using the mixed data, the performance of DeepFlu dropped for identifying the susceptible to H1N1 or H3N2, suggesting there may exist distinct differences in susceptibility to different IAV strains.

There are some limitations in this study. First, limited influenza gene expression data are available. Although DeepFlu excels even with limited gene expression data, the deep learning model may improve further with more training data. Second, it is unknown if the subjects were immunologically naïve to IAV. Such conditions may need to be controlled for a more appropriate comparison or validation. Third, the biological mechanisms behind IAV susceptibility or immunity remain unclear. DeepFlu with full features, which does not rely on specific gene sets, may require a subtle approach to further study the mechanisms behind IAV susceptibility or immunity.

## 5. Conclusion

Gene expression can portend IAV susceptibility and deep learning can catch the expression signals to identify people at risk for the flu and alarm them in advance to reduce hospitalization and mortality caused by IAV.

## Acknowledgements

This study was supported by Ministry of Science and Technology, R.O.C. under MOST-108-2221-E-019-052. The funders had no role in study design, data collection and analysis, decision to publish, or preparation of the manuscript.

## Supplementary material

Supplementary material associated with this article can be found, in the online version.

